# Optimisation and validation of a sensitive bioanalytical method for niclosamide

**DOI:** 10.1101/2021.01.13.426426

**Authors:** Usman Arshad, Henry Pertinez, Helen Box, Lee Tatham, Rajith KR Rajoli, Megan Neary, Joanne Sharp, Anthony Valentijn, James Hobson, Catherine Unsworth, Andrew Dwyer, Alison Savage, Tom O Mcdonald, Steve P Rannard, Paul Curley, Andrew Owen

## Abstract

The SARS-CoV-2 pandemic has spread at an unprecedented rate, and repurposing opportunities have been intensively studied with only limited success to date. If successful, repurposing will allow interventions to become more rapidly available than development of new chemical entities. Niclosamide has been proposed as a candidate for repurposing for SARS-CoV-2 based upon the observation that it is amongst the most potent antiviral molecules evaluated *in vitro*. To investigate the pharmacokinetics of niclosamide, reliable, reproducible and sensitive bioanalytical assays are required. Here, a liquid chromatography tandem mass spectrometry assay is presented which was linear from 31.25-2000 ng/mL (high dynamic range) and 0.78-100 ng/mL (low dynamic range). Accuracy and precision ranged between 97.2% and 112.5%, 100.4% and 110.0%, respectively. The presented assay should have utility in preclinical evaluation of the exposure-response relationship and may be adapted for later evaluation of niclosamide in clinical trials.

## Introduction

Niclosamide (5-chloro-N-(2-chloro-4-nitrophenyl)-2-hydroxybenzamide) is an Food and Drug Administration (FDA) approved anthelmintic drug that is listed as an Essential Medicine by the WHO [1]. It has been in use since the 1960s, primarily for treating tapeworm infections [2]. Niclosamide has been proposed as a candidate for repurposing for SARS-CoV-2 based upon the observation that it is amongst the most potent molecules evaluated *in vitro* with VeroE6 cells [3]. At the time of writing, no studies have assessed the efficacy of niclosamide by any route of administration in an animal model. However, 10 studies are currently listed on clinicaltrials.gov describing trials of either oral or inhaled niclosamide formulations (www.clinicaltrials.org).

The putative mechanism of action for niclosamide is not currently fully understood for SARS-CoV-2 but preliminary data have started to emerge within the preprint literature [4] A thorough understanding of the mechanism of action is critical to rationalize whether *in vitro* observations can be expected to lead to effects *in vivo.* Successful development of antiviral interventions for SARS-CoV-2 will also require a robust assessment of the pharmacokinetic-pharmacodynamic relationship in preclinical models and patients, and many groups have called for better application of the principles of antiviral pharmacology during candidate selection for COVID-19 [3, 5, 6]. To better understand the exposure-response relationship, robust and validated bioanalytical methods are required to accurately determine the pharmacokinetics in preclinical and clinical studies.

Many existing methods for the analysis of niclosamide rely on HPLC systems [7–9]. Such methods are susceptible to interferences by other chemicals and are limited by poor sensitivity. While LC/MS MS methods for the detection of niclosamide have been developed; many are specific to matrices not representative for *in vivo* samples (e.g. water or cell culture media), or lack the dynamic range required for application to pharmacokinetic (PK) studies [10–12]. The potential application of niclosamide to SARS-CoV-2 therapy presents a bioanalytical challenge because of the paucity of data available publicly for the pharmacokinetics following administration to humans [13]. As such, development of a rapid, sensitive and versatile method to determine niclosamide concentrations in biological matrices is warranted.

The assay presented here was developed and validated in accordance with FDA guidelines [14]. Criteria such as linearity, accuracy, precision, selectivity (ensuring detection of the analyte and not an endogenous compound within the sample matrix) and recovery were assessed. The presented assay has application in pre-clinical evaluation as a SARS-CoV-2 antiviral intervention and may be adaptable to human matrices for future clinical evaluation if warranted.

## Materials and Methods

### Materials

Niclosamide and the internal standard (IS) tizoxanide were purchased from Stratech Scientific Ltd (Cambridge, UK). Drug free Sprague Dawley plasma with lithium heparin was purchased from VWR International (PA, USA). LCMS grade acetonitrile (ACN) was purchased from Fisher Scientific (MA, USA). All other consumables were purchased from Merck Life Science UK LTD (Gillingham, UK) and were of LC-MS grade.

### Tuning and Calibration

Detection of niclosamide was conducted using a QTRAP 6500+ Mass Spectrometer coupled with an Exion AD liquid chromatography system (SCIEX, UK). Tuning was performed using direct infusion (10 μL/min) of a 10ng/mL stock of niclosamide with H_2_O:methanol (50:50). Ionisation was achieved via heated electron spray ionisation in the negative mode. The multiple reaction monitoring mode was selected for quantification of thea analytes, for which the precursors to production ion transitions were as follows: niclosamide 324.913→170.8 and 324.913→288.8. MS parameter voltage settings (DP, EP, CE, and CXP) were optimised to achieve the highest signal intensity.

### Chromatographic separation

Chromatographic separation was achieved using a multistep gradient (Table 1) with a Kinetex^®^ C18 column (100×2mm, 2.6 μM) obtained from Phenomenex (Macclesfield, UK). Mobile phases A and B were H_2_O + 0.05% formic acid and ACN + 0.05% formic acid, respectively. The assay was conducted over 3.5 minutes at a flow rate of 400 μl/min, column temperature 25°C. Data acquisition and chromatography analysis were carried out using the Analyst Software.

**Table 1.** The operating chromatographic conditions

| Time (minutes) | Mobile Phase A (%) | Mobile Phase B (%) |
| --- | --- | --- |
|  | H<sub>2</sub>O 0.05% formic acid | ACN 0.05% formic acid |
| 0.0 | 95 | 5 |
| 0.5 | 95 | 5 |
| 1 | 20 | 80 |
| 2.5 | 5 | 95 |
| 3.0 | 5 | 95 |
| 3.01 | 95 | 5 |
| 3.5 | 95 | 5 |

### Preparation of standards and controls

Stock solutions of niclosamide and the IS tizoxanide were prepared at a concentration of 1mg/mL in ACN and stored in glass vials at 4°C. Working standards of niclosamide were prepared in rat plasma via serial dilution ranging from 31.25 to 2000 ng/mL for the high dynamic range or 0.78 to 100 ng/mL for the low dynamic range. Quality control samples (QC) of 75, 750 and 1500ng/mL or 4, 40 and 80 ng/mL were prepared, respectively. The IS working solution (10 ng/mL) was prepared by diluting its stock solution with ACN.

### Extraction procedures

Extraction was performed using a protein precipitation method. For the high dynamic range, a total volume of 100 μl of sample, standard or QC was transferred to glass vials where 500 μL of 100% ACN containing 10 ng/mL of IS was added. Following mixing, samples were centrifuged at 3500g for 5 minutes at room temperature. Thereafter, 50 μl of the supernatant was directly transferred into 200 μl chromatography vials. A further 50 μl of H_2_0 was added and 1 μl of each sample was injected into the LC-MS/MS system for analysis.

Similarly, for the low dynamic range a total volume of 100 μl of sample, standard or QC was transferred to glass vials where 500 μL of 100% ACN containing 10 ng/mL of IS was added. Following mixing, samples were centrifuged at 3500g for 5 minutes at room temperature. 500 μL of supernatant fraction was transferred to a fresh glass vial and evaporated to dryness under a gentle stream of nitrogen. The pelleted residue was reconstituted with 100 μL solution of H_2_O:ACN (50:50) and vortexed. Subsequently, 50 μl of the sample was then transferred into 200 μl chromatography vials. Thereafter 1 μl of each sample was injected into the LC-MS/MS system for analysis.

### Assay Validation

The assay was validated according to the FDA guidelines for the development of bioanalytical assays [14] The following criteria were assessed: linearity, recovery, selectivity, accuracy, precision and inter-assay as well as intra-assay variability.

### Linearity

Linearity was assessed by three independent preparations of a calibration standard curve for each analyte. The calibration curve was obtained by plotting the ratio of peak area of analyte to the peak area of IS. Maximum allowed deviation of standards was set at 15% from the expected value, excluding the lower limit of quantification (LLOQ) where deviation was set at no more than 20%.

### Recovery

Recovery experiments were performed by comparing the results for extracted samples of each compound at three concentrations (same as high, medium and low QCs) with non-extracted standards that were taken to represent 100% recovery.

### Selectivity

The degree of interference from the matrix (due to potential interfering substances including endogenous matrix components, metabolites and decomposition products) was assessed via comparison of extracted blank samples with the lowest point of the standard curve, the LLOQ. The LLOQ was a minimum of five-times greater than the background signal.

### Accuracy & precision

The intra-day and inter-day precision and accuracy of the method were determined with QC samples at three different concentrations (N=3) on the same day and on three different days. The calculated mean concentration relative to the nominal concentration was used to express accuracy (% variability of accuracy = error/stated value*100). The relative standard deviation was calculated from the QC values and used to estimate the precision (%variation of precision = standard deviation/mean assay value*100). Acceptable variation for accuracy and precision was set at 15% and at 20% for the lower concentrations.

## Results

### Method development

The optimised global settings were: Ion spray voltage −4500V, Curtain gas 25, source temperature 450°C and Ion source gas (GS1/GS2) 50.00. MS parameter voltage settings for niclosamide were fine-tuned for maximum sensitivity, and parent ion transitions (MRM mode) were selected to afford the best response for the spectrum analysis as shown in the table below.

### Linearity

The standard curves were linear in the concentration range between 31.25-2000 ng/mL and 0.78-100 ng/mL (Fig 1). The average correlations of calibration curve were found to be acceptable (Pearson’s coefficient of determination R^2^>0.99). The peak area ratio (analyte to IS; variation of IS was <15% in each run) was proportional to the stated concentration ranges.

**Figure 1.**
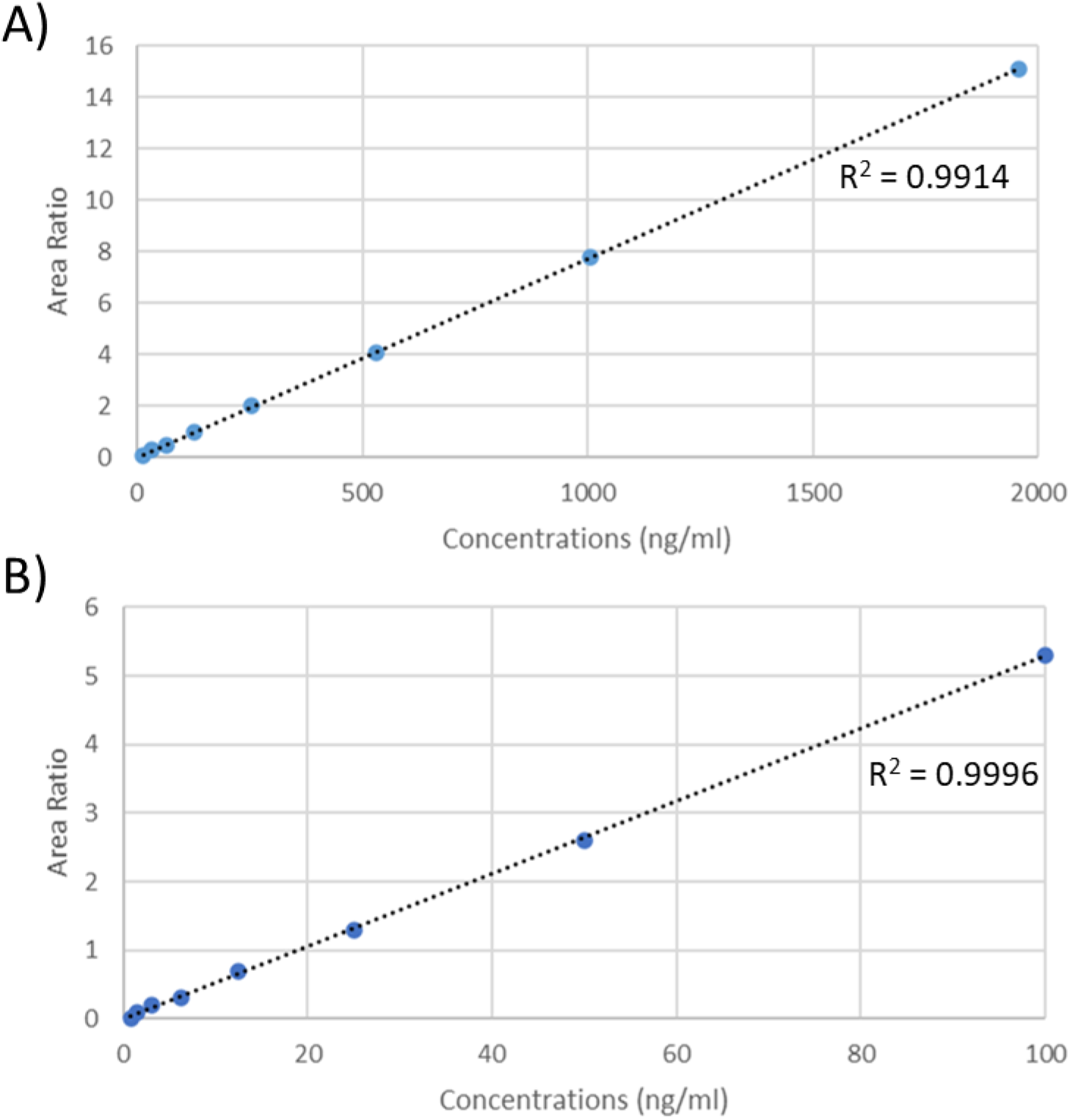
Linearity of the bioanalytical method. A) shows an example of the linearity using the high concentration range (31.25-2000ng/mL). B) shows an example of the linearity using the low concentration range (0.78-100ng/mL). R^2^ values for the line of best fit are also shown.

### Extraction Recovery

The recovery was evaluated using three replicates of QC samples at three concentration levels (the same concentrations as QC sample) in rat plasma. The results (Fig 2) showed that the recoveries at all three concentrations within both methods to be >70%. The mean recovery (across the 3 QCs) in the high dynamic range (Fig 2A) were 75% (standard deviation 3.17) for the low QC, 75% (standard deviation 2.95) for the mid QC, and 74% (standard deviation 1.84) for the high QC. Recoveries for the low dynamic range (Fig 2B) were 97% (standard deviation 2.05) for the low QC, 97% (standard deviation 1.59) for the mid QC, and 97% (standard deviation 1.33) for the high QC.

**Figure 2.**
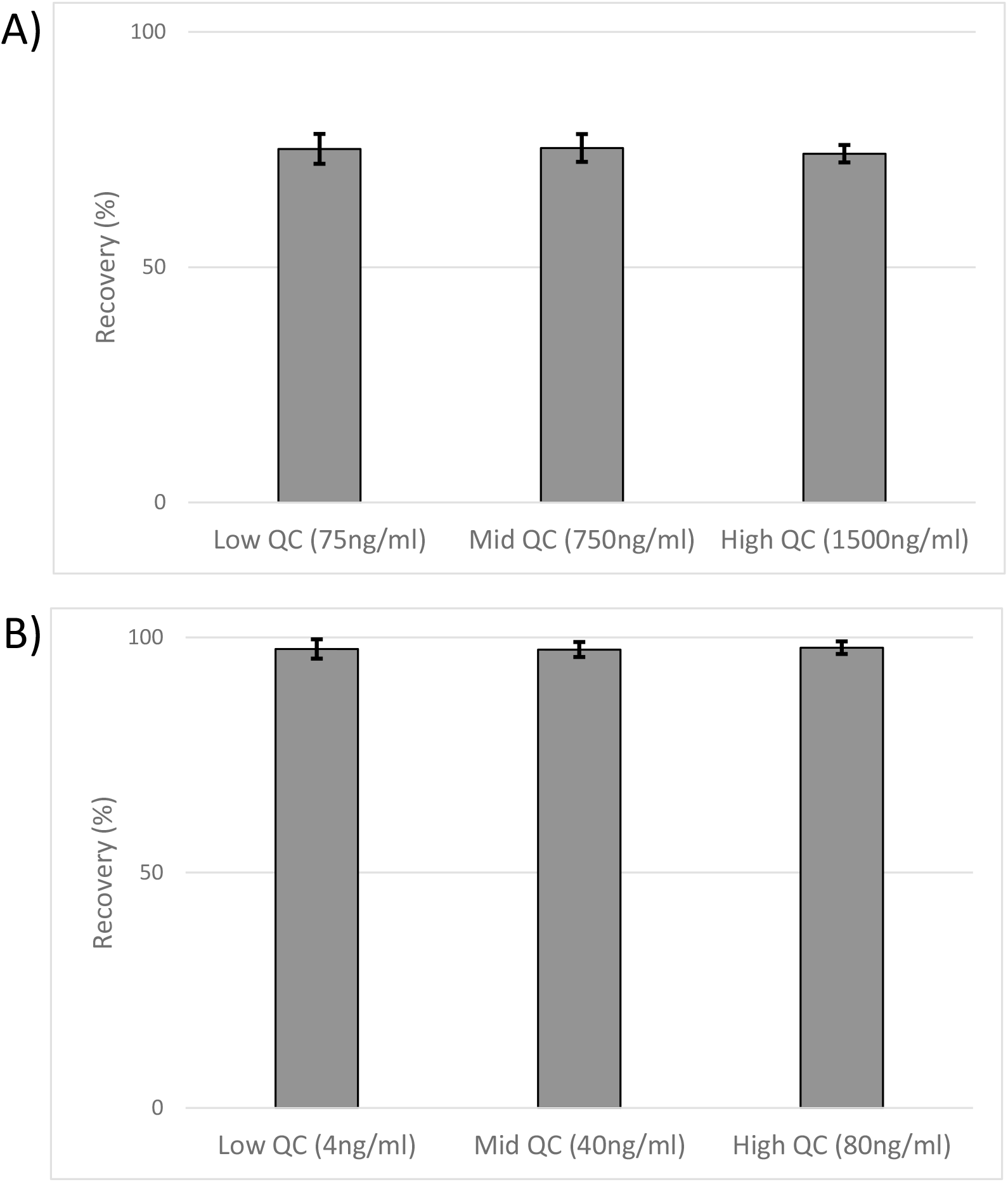
Niclosamide recovery. A) shows the percentage recovery for high dynamic range and B) shows the percentage recovery for low dynamic range. Data is show percentage of unextracted standards ± percentage standard deviation.

### Selectivity

The matrix effect of plasma was examined by comparing extracted blank plasma to extracted plasma spiked with tizoxanide. For the high dynamic range figure 3A shows the chromatogram produced by the extracted blank. There was no visible peak (area NA) at the retention time of niclosamide (2.5 minutes). FDA guidelines require the lower limit of quantification produce a peak area of at least five-fold greater than that observed in the blank matrix. Figure 3B shows the peak produced from the lower limit of quantification (31.25 ng/mL). The peak area is 2960, which complies with FDA guidelines. Figure 3C shows the peak (354170) produced by the highest standard (2000 ng/mL). The bottom panels of figure 3 (D, E, F) for the low dynamic range also complied with FDA guidelines. Peak areas were as follows, 26518, 207766 and 25351511, for the extracted blank, the lower limit of quantification (0.78 ng/mL) and the highest standard (100 ng/mL), respectively. Additionally, the signal produced by the IS showed no interference with niclosamide in either methods.

**Figure 3.**
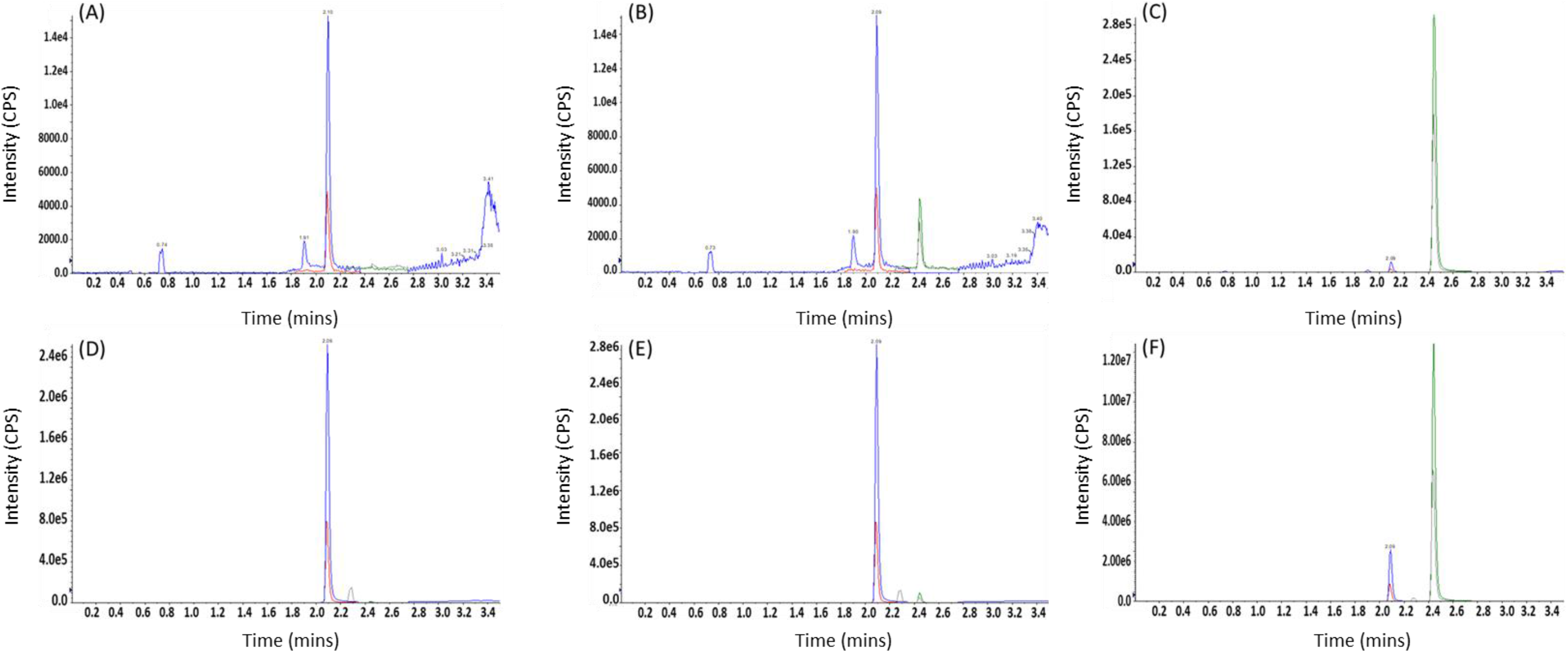
shows representative chromatograms from the high dynamic range method for the blank plasma (A), lower limit of quantification (31.25ng/mL) (B) and the highest standard (2000ng/mL) (C). Representative chromatograms for the low dynamic range method are shown for the blank plasma (D), lower limit of quantification (0.78ng/mL) (E) and the highest standard (100ng/mL) (F). The peak produced by niclosamide had a retention time of 2.5 minutes and the peak produced by the IS tizoxanide had a retention time 2.09 minutes.

### Accuracy & precision

The variability within and between assays was calculated to demonstrate that the accuracy and precision were maintained across repetitions of the assay. Table 3 shows the variance of accuracy and precision calculated from mean values of three repetitions of the assay (low QC, medium QC and high QC) either from the same experiment (intra-day) or an average of three separate experiments (inter-day). These data demonstrate that the precision and accuracy values were in the acceptance range (<15%). The intra and inter-day precision was ≤3.42% and ≤10% for niclosamide. The intra and inter-day accuracy was ≤12.5% and ≤2.9% for niclosamide.

**Table 2.**
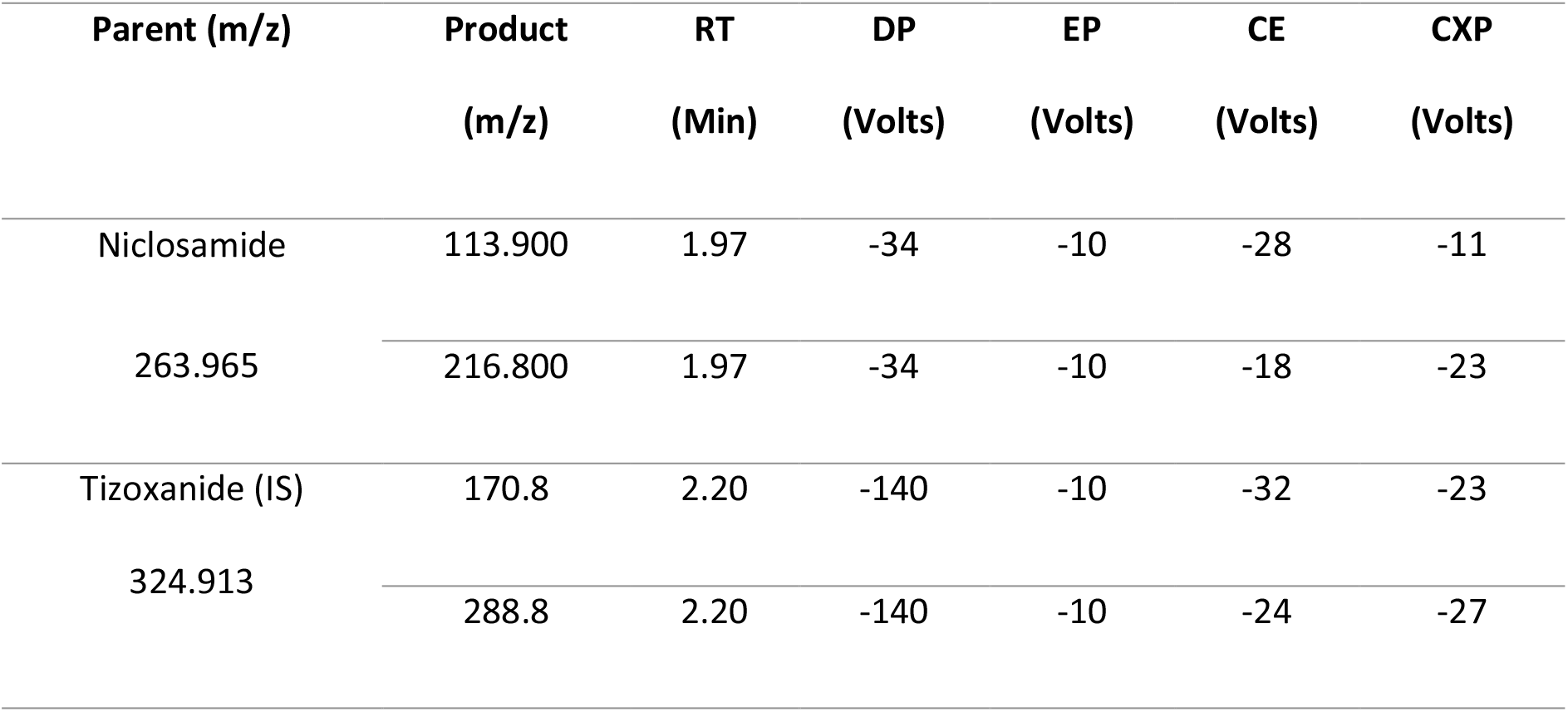
shows the final source parameters for used for the detection of niclosamide and tizoxanide (IS).

**Table 3.** Intra-day and Inter-day accuracy and precision for niclosamide in rat plasma (n=3)

|  |  | Intra-day |  |  | Inter-day |  |  |
| --- | --- | --- | --- | --- | --- | --- | --- |
| | | Average ± SD | | | Average ± SD | | |
|  |  | (ng/mL) | Variance of accuracy (%) | Variance of precision (%) | (ng/mL) | Variance of accuracy (%) | Variance of precision (%) |
| **Niclosamide High Dynamic Range** | High (1500 ng/mL) | 1518.0 ± 28.2 | 1.2 | 1.86 | 1514.2 ± 3.33 | 0.94 | 0.22 |
| | Medium (750ng/mL) | 741.5 ± 14.8 | -1.12 | 1.9 | 751.2 ± 8.4 | 0.2 | 1.12 |
| | Low (75ng/mL) | 72.8 ± 2.0 | -2.8 | 2.74 | 75.1 ± 1.9 | 0.1 | 2.5 |
| **Niclosamide Low Dynamic Range** | High (80 ng/mL) | 82.2 ± 2.8 | 2.75 | 3.42 | 78.3 ± 5.6 | -2.1 | 7.1 |
| | Medium (40 ng/mL) | 44.9 ± 0.1 | 12.3 | 0.4 | 41.1 ± 3.9 | 2.9 | 9.6 |
| | Low (4ng/mL) | 4.5 ± 0.1 | 12.5 | 2.7 | 4.1 ± 0.4 | 1.9 | 10.0 |

## Discussion

The SARS-CoV-2 pandemic has spread at an unprecedented rate, and repurposing opportunities have been intensively studied with only limited success to date. If successful, this approach will allow interventions to become rapidly available but a thorough understanding of the exposure-response relationship is required to improve the chances of success [3]. Robust bioanalytical methods underpin this understanding, providing a basis for translation of candidates to clinical evaluation. However, ultimate utility can only be confirmed in large and appropriately controlled randomized clinical trials, and drugs should not be used until such data are rigorously assessed with approval by appropriate agencies.

The assay presented here was developed for use in preclinical assessment of niclosamide pharmacokinetics in rats. Validation was conducted in plasma satisfying FDA bioanalytical method development guidelines, demonstrating good accuracy, precision and linearity. This assay represents an excellent starting point for researchers seeking to develop niclosamide assays. It should be noted that a very low signal was produced by blank samples, and the lower limit of detection was not fully evaluated in this study. Therefore, the true limit of the assay may be lower than the range reported here and this could be further optimized if required. The presented assay is currently being adapted to quantify niclosamide from plasma from other preclinical species and human plasma. Additional work is also underway to adapt the assay for quantification in other tissues such as lung and nasal turbinate. However, due to the more complex nature of tissue homogenates considerable additional optimization may be required. Tissue homogenate has a different protein composition compared to plasma and common proteins may be in different abundance. Niclosamide has been shown to bind to various endogenous proteins including hemoglobin, human serum albumin, and globulin demonstrating the importance of assessing any potential change in matrix when adapting this assay [15, 16].

Examples of contemporary assays demonstrate niclosamide detection in a variety of matrices. The sensitivity typically falls in the range of approximately 1-40 ng/mL with an upper limit of 1000-8000 ng/mL [7, 11, 13]. Although assays have been reported with lower sensitivity and higher detection limits, these assays are limited by the dynamic range. For example, Wang *et al* describe a method with a range of 0.02-12.5 ng/mL for *in vitro* applications [10]. While the lower limit of quantification is better than the assay presented here, the upper limit is not suitable for application to *in vivo* pharmacokinetic evaluation. Additionally, cell culture media is a considerably less complex matrix than plasma.

In summary, the optimization of a robust assay is described, providing a simple and sensitive LC-MS/MS assay for application in preclinical assessments of niclosamide. The final assay conformed to FDA bioanalytical development guidelines and may be used to further the ongoing investigations into the utility of niclosamide against SARS-CoV-2, irrespective of formulation or route of administration.

## Conflicts of interest statement

AO and SR have received research funding from AstraZeneca and ViiV and consultancies from Gilead; AO has additionally received funding from Merck and Janssen and consultancies from ViiV and Merck not related to the current paper. No other conflicts are declared by the authors.

## Funding

This work was funded by UKRI using funding repositioned from EP/R024804/1 as part of the UK emergency response to COVID-19. The authors also acknowledge research funding from EPSRC (EP/S012265/1), NIH (R01AI134091; R24AI118397), European Commission (761104) and Unitaid (project LONGEVITY).

